# Alpha-frequency stimulation strengthens coupling between temporal fluctuations in alpha oscillation power and default mode network connectivity

**DOI:** 10.1101/2025.01.27.635137

**Authors:** Yijia Ma, Joshua A. Brown, Chaowen Chen, Mingzhou Ding, Wei Wu, Wen Li

## Abstract

Alpha (8-12 Hz) oscillations and default mode network (DMN) activity dominate the brain’s intrinsic activity in the temporal and spatial domains, respectively. They are thought to play crucial roles in the spatiotemporal organization of the complex brain system. Relatedly, both have been implicated, often concurrently, in diverse neuropsychiatric disorders, with accruing electroencephalogram/magnetoencephalogram (EEG/MEG) and functional magnetic resonance imaging (fMRI) data linking these two neural activities both at rest and during key cognitive operations. Prominent theories and extant findings thus converge to suggest a mechanistic relationship between alpha oscillations and the DMN. Here, we leveraged simultaneous EEG-fMRI data acquired before and after alpha-frequency transcranial alternating current stimulation (α-tACS) and observed that α-tACS tightened the dynamic coupling between spontaneous fluctuations in alpha power and DMN connectivity (especially, in the posterior DMN, between the posterior cingulate cortex and the bilateral angular gyrus). In comparison, no significant changes were observed for temporal correlations between power in other oscillatory frequencies and connectivity in other major networks. These results thus suggest an inherent coupling between alpha and DMN activity in humans. Importantly, these findings highlight the efficacy of α-tACS in regulating the DMN, a clinically significant network that is challenging to target directly with non-invasive methods.

**Significance Statement:** Alpha (8-12 Hz) oscillations and the default mode network (DMN) represent two major intrinsic activities of the brain. Prominent theories and extant findings converge to suggest a mechanistic relationship between alpha oscillations and the DMN. Combining simultaneous electroencephalogram-functional-magnetic-resonance imaging (EEG-fMRI) with alpha-frequency transcranial alternating current stimulation (α-tACS), we demonstrated tightened coupling between alpha oscillations and DMN connectivity. These results lend credence to an inherent alpha-DMN link. Given DMN dysfunctions in multiple major neuropsychiatric conditions, the findings also highlight potential utility of α-tACS in clinical interventions by regulating the DMN.

---

Research using resting-state (RS) electroencephalography (EEG), magnetoencephalography (MEG), and functional magnetic resonance imaging (fMRI) recordings has reliably depicted the temporal and spatial landscapes of intrinsic activities in the human brain. Temporally, as revealed by EEG/MEG, the resting brain is dominated by oscillatory activities in the range of 8-12 Hz (known as alpha oscillations), representing the main rhythm of intrinsic neural synchronization [1, 2]. Spatially, as demonstrated by fMRI, the resting brain is characterized by widely distributed coactivation anchored in a midline core comprising the posterior cingulate cortex (PCC) and the medial prefrontal cortex (mPFC), along with the left and right angular gyri (AG), collectively forming the default mode network (DMN) [3, 4].

Both the DMN and alpha oscillations underpin the dynamic organization of the brain. Through long-range synchrony and coactivation, alpha oscillations and the DMN can temporally and spatially orchestrate neural processes on meso- and macro-scopic levels, constituting pivotal architectures of the brain’s intrinsic organization [5–8]. As such, alpha oscillations and the DMN are both strongly implicated in fundamental cognitive operations, including consciousness, awareness, and vigilance [4, 9]. Similarly, their dysfunctions have been linked to the pathophysiology of major neuropsychiatric disorders, including Alzheimer’s disease, schizophrenia, and post-traumatic stress disorder (PTSD), representing fundamental, transdiagnostic pathological underpinnings [10–16].

Akin to these prominent parallels between them, growing evidence suggests that alpha oscillations and the DMN are inherently and mechanistically associated [17, 18]. Research combining high-density EEG/MEG with fMRI or source localization indicates that alpha oscillations represent the primary neural synchrony linking the core hubs (mPFC and PCC) of the DMN [18–21]. In PTSD, this alpha-oscillatory connectivity between the DMN midline hubs is deficient, underscoring the joint alpha-DMN pathology [22]. In addition, alpha oscillations and DMN activity are highly dynamic, fluctuating (“waxing and waning”) spontaneously over time [23, 24]. Research has leveraged these temporal dynamics to directly link alpha and DMN activities: simultaneous EEG-fMRI recordings have repeatedly revealed that the spontaneous fluctuations in alpha oscillations and DMN activity are temporally coupled [19, 25–28], thereby highlighting a direct linkage.

Recently, using transcranial alternating current stimulation (tACS) at the alpha frequency to augment alpha oscillations, research has further demonstrated that alpha enhancement leads to the upregulation of DMN connectivity and activation [29, 30]. These findings thus highlight a mechanistic alpha-DMN association, with shared neural underpinnings that are responsive to alpha-frequency tACS (α-tACS). By that token, we hypothesized that by upregulating the shared neural underpinnings, α-tACS would not only result in the concurrent enhancement of alpha and DMN activity but also tighten their temporal coupling. Evidence for this hypothesis would lend further credence to the potentially mechanistic linkage between alpha oscillations and the DMN. Importantly, it would constitute a mechanistic foundation for non-invasive brain stimulation such as α-tACS to upregulate these two systems, which both figure critically in mental functioning and its various disorders.

Therefore, this study sought to directly examine the effect of α-tACS on the temporal coupling of spontaneous alpha and DMN fluctuations. Towards that end, we reanalyzed data from [29], where RS EEG and fMRI were simultaneously recorded before and after 20-minutes of occipitoparietal α-tACS. Extracting the time series of alpha power and DMN connectivity and comparing their correlation (reflecting the temporal coupling) before and after stimulation, we tested the hypothesis that the coupling of alpha and DMN fluctuations would increase after α-tACS.

## Methods

### Participants

Data from a prior study [29] were re-analyzed to test the hypotheses. Thirty-two healthy participants (17 female; age 20.9 +/- 3.5 years), who had simultaneous EEG-fMRI data, were included in the current study. Participants had no prior history of neurological or psychiatric disorders or use of psychotropic medications at time of recruitment. Participants were randomly assigned to receive either active tACS stimulation (*n* = 16) or sham stimulation (*n* = 16). Groups did not differ in age or gender distribution (*p*’s > 0.3).

### Experimental Procedures

The experiment consisted of three phases: pre-stimulation RS recordings, tACS/sham stimulation, and post-stimulation RS recordings (**Figure 1A**). In both pre- and post-stimulation phases, participants underwent two successive 5-minute (10 minutes in total) simultaneous EEG-fMRI scans (with eyes open and fixated on a central crosshair). α-tACS was administered for 20 minutes, with a ±2 mA sinusoidal current oscillating at 10 Hz using an MR-compatible High-Definition (HD) tACS system (Soterix Medical New York, NY, USA). A 4 × 1 montage over midline occipitoparietal sites (with 4 surrounding + 1 central electrodes forming a closed circuit) was selected to maximally target the primary cortical source of alpha oscillations— occipitoparietal cortex [1, 2, 31]. More details are reported in [29].

**Figure 1.**
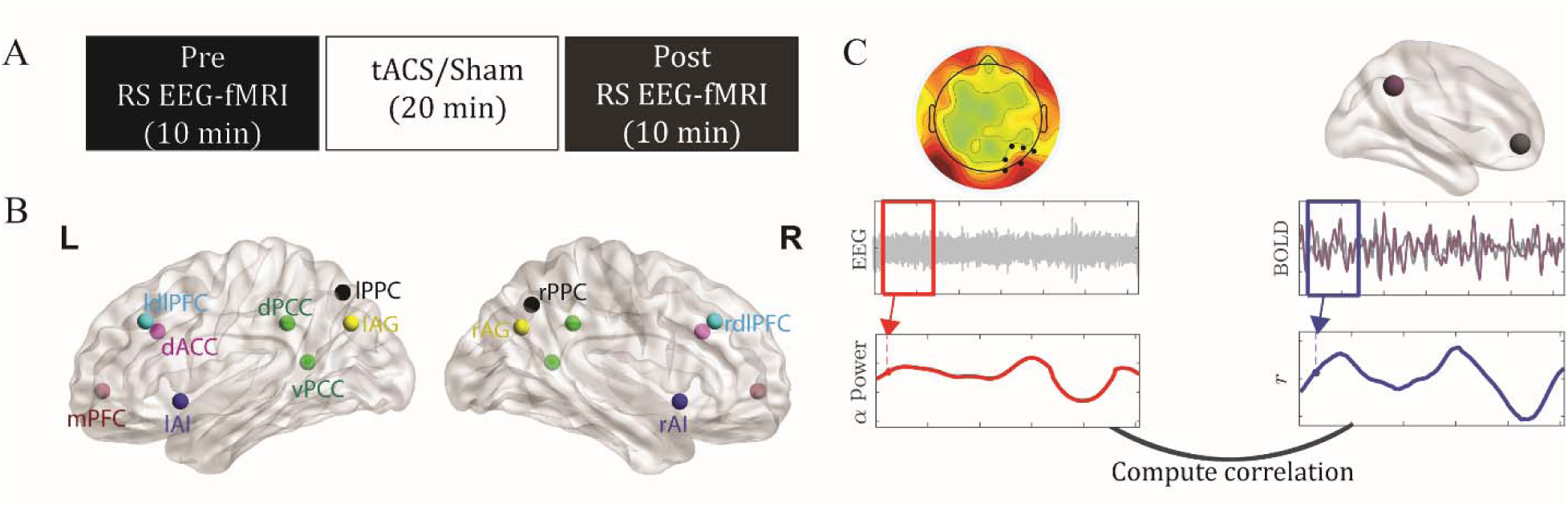
Experimental paradigam (A), Regions of interest (B) and dynamic analysis pipeline (C). (**A**) Experimental design. Participants underwent simultaneous RS EEG-fMRI recordings before and after tACS/Sham stimulation. Each recoding session lasts ten minutes. (B) Regions of interest; Abbreviations: (l/r) AI = (left/right) anterior insula; dACC, dorsal anterior cingulate cortex; (l/r) dlPFC = (left/right) dorsolateral prefrontal cortex; (d/v) PCC = (dorsal/ventral) posterior cingulate cortex; (l/r) PPC = (left/right) posterior parietal cortex; mPFC = medial prefrontal cortex; (l/r) AG = (left/right) angular gyrus. (**C**) Analysis pipeline illustrating the methods used (see text for more details).

### fMRI acquisition and preprocessing

fMRI data were acquired in a 3T Siemens Prisma scanner with a 64-channel head coil and axial acquisition. Imaging parameters included TR/TE: 1,800/22.40 ms; slice thickness 1.8 mm; gap 0.45 mm; in-plane resolution/voxel size 1.8 × 1.8 mm; multiband acceleration factor = 2; GeneRalized Autocalibrating Partial Parallel Acquisition (GRAPPA) acceleration factor = 2. A high resolution (0.9 × 0.9 × 0.9 mm^3^) three-dimensional Magnetization Prepared Rapid Acquisition Gradient Echo (3D-MPRAGE) T1 scan was also acquired. Following discarding dummy scans (the first 9 volumes of each block), data of the two blocks were concatenated.

Imaging data were preprocessed using SPM12, including slice-time correction, spatial realignment, and normalization using Diffeomorphic Anatomical Registration Through Exponentiated Lie algebra (DARTEL). To further remove artifacts potentially contributing to spurious RS activity variance [32], we implemented additional preprocessing using the Data Processing Assistant for Resting-State fMRI (DPARSFA) toolbox: 1) mean centering and whitening of timeseries; 2) temporal bandpass (0.01 to 0.08 Hz) filtering; and 3) general linear modeling to partial out head motion with 24 nuisance variables (six head motion parameters each from the current and previous scan and their squared values).

#### Regions of interest (ROIs)

ROIs were selected from the hubs of three major intrinsic connectivity networks (ICNs), namely, the default mode network (DMN), executive control network (ECN), and salience network (SN). The ECN and SN were included as control networks to elucidate whether alpha oscillations are specifically coupled with the DMN.

We defined these ROIs primarily based on the 90 functional ROI Willard atlas [33]. DMN ROIs included the ventromedial prefrontal cortex (mPFC), the posterior cingulate cortex (PCC), and the left and right angular gyrus (l/r AG). As in the previous study [29], given that the PCC hub consists of two functionally-dissociable, ventral and dorsal, subdivisions (with the ventral PCC/vPCC primarily linked with DMN nodes and the dorsal PCC/dPCC with extensive extra-DMN connections [4, 34]), we defined the vPCC and dPCC individually from the Brainnetome Atlas [35]. ECN ROIs included the left and right dorsolateral prefrontal cortex (l/r dlPFC) and posterior parietal cortex (l/r PPC). SN ROIs included the dorsal anterior cingulate cortex (dACC) and the left and right anterior insula (l/r AI), See Figure 1(B).

### EEG acquisition and preprocessing

EEG data were recorded using an MR-compatible 64-channel system (Brain Products GmbH, Germany). An additional electrode was placed on the participant’s back to record electrocardiogram (ECG), allowing for the removal of cardiac artifacts from the EEG data. Detailed information about data acquisition, preprocessing, and artifact removal is provided in [29]. The cleaned EEG data were segmented into epochs of 1.8 s (i.e., the TR length) centered on the onset of each fMRI scan (TR).

Oscillatory power was computed at individual channels for each epoch using the multitaper spectral estimation technique with 3 tapers [36, 37]. Alpha (8–12 Hz) power was extracted from the right occipitoparietal site (collapsed across P4, P6, P8, PO4, PO8 and O2) which demonstrated the strongest alpha increase following α-tACS [29] (**Figure 1C**). These values were then normalized by the mean power for the global spectrum (1–40 Hz). To determine alpha-frequency-specific effects, power was similarly derived for the neighboring frequencies, theta (4-7 Hz) and low beta (13-17) bands, and submitted to similar analyses.

### Dynamic Analysis

To examine the temporal coupling between alpha oscillations and DMN connectivity, we measured the covariation of fluctuations of these two activities over time. We implemented a tapered sliding windows approach as commonly used in previous studies [38–40]. We used a Gaussian window, with smoothing kernel/FWHM = 27.78 epochs and a length of 108 seconds (60 epochs), resulting in a frequency threshold of 0.01 Hz. This threshold, corresponding to the lower bound of the RS BOLD activity, has been commonly used [41–46]. To ensure sufficient temporal resolution, we selected sliding increments of 1.8 s (1 TR), which resulted in 247 windows (i.e., time points) with an overlap of 106.2 s (59 TR) between neighboring windows [17, 47, 48].

We extracted BOLD signals from every ROI within each Gaussian window and computed the functional connectivity between the ROIs for each window. We convolved EEG power per epoch with the Gaussian window to generate the EEG power timeseries. We measured the covariation between fluctuations of fMRI connectivity and oscillatory power by extracting Pearson correlation coefficients between the fMRI functional connectivity timeseries and EEG power timeseries. Following previous work [49], we avoided convolving the EEG power timeseries with a hemodynamic response function such that its amplitude values would be preserved without potential distortions. To assess whether the inherent time resolutions between imaging modalities affected the results, we performed validation analyses varying the offset between EEG and fMRI. **Figure 1** illustrates the procedures of this analysis. All computations were performed using in-house MATLAB scripts, relying solely on built-in functions, except for notBoxPlot (https://github.com/raacampbell/notBoxPlot/) for visualization and select EEGLAB functions [50] for EEG data processing.

### Statistical analysis

To test our hypothesis that tACS would enhance the dynamic coupling between alpha oscillations and DMN connectivity, we submitted the Pearson correlation coefficients (after Fisher Z transformation) to double contrasts: Post – Pre_(Active-Sham)_. The double contrasts were corrected for multiple comparisons across the DMN hubs, based on FDR *p* < .05. We followed up the significant double contrasts with simple contrasts to unpack the effects. As these simple contrasts were restricted to (and hence, protected by) the significant double contrasts, we did not apply further corrections (to prevent over-correction that could increase Type II errors). For the same reason, we also used one-tailed *p* < .05 for the simple contrasts.

## Results

### α-tACS enhanced alpha-DMN coupling

Our double contrasts indicated an increase in the coupling between alpha power and vPCC-lAG connectivity timeseries in the Active (versus Sham) group after stimulation (*t* = 3.14, FDR *p* = 0.038; **Figure 2**). Specifically, the Active group showed an increase (*t* = 2.57, *p* = 0.011, one-tailed), while the Sham group showed a decrease (*t* = -1.79, *p* = 0.046, one-tailed) in this coupling. Our double contrasts also showed an increase in the coupling between alpha power and vPCC-rAG connectivity timeseries in the Active (versus Sham) group (*t* = 2.76, FDR *p* = 0.048) after stimulation. Specifically, the Active group exhibited an increase (*t* = 1.82, *p* = 0.045, one-tailed) compared to the Sham group with a decrease (*t* = -2.16, *p* = 0.023, one-tailed) in this coupling. Nuances of such effects are illustrated in **Figure 3**, based on data from a representative participant in the Active group. Finally, our control analyses below further ruled out a possible confound of global temporal association between EEG-fMRI-connectivity (**Figure 5**).

**Figure 2.**
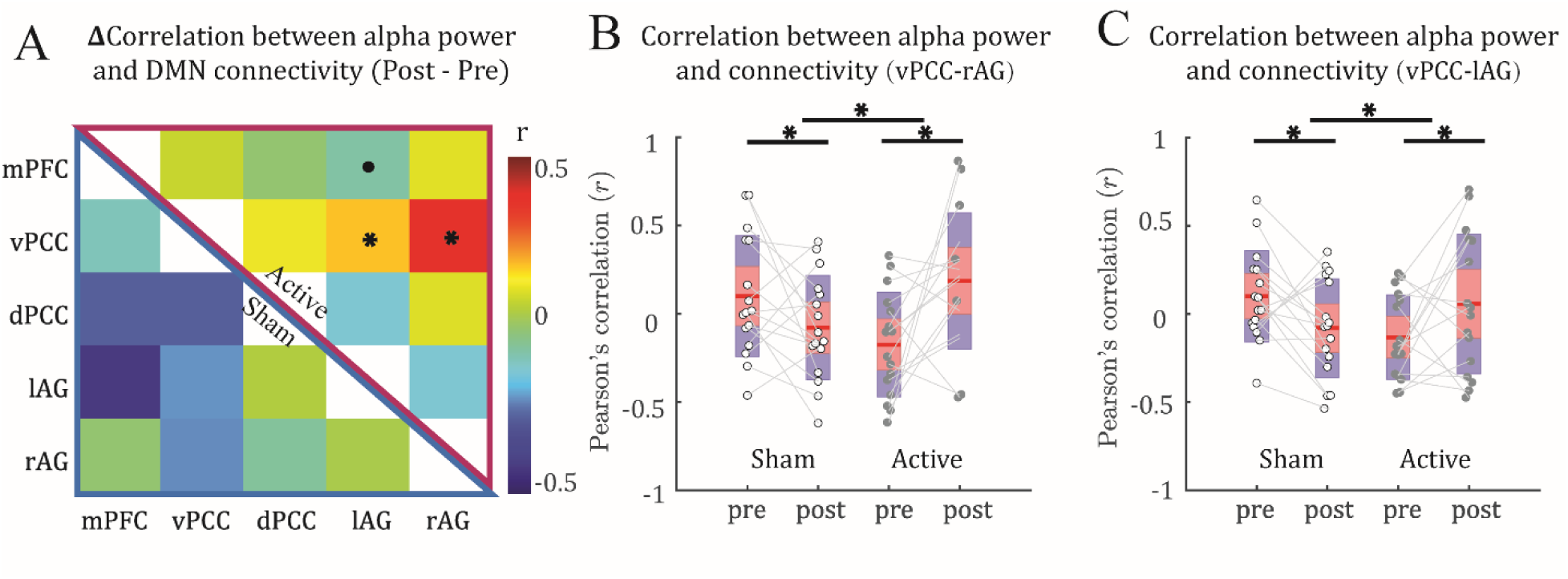
*α*-tACS strengthened dynamic coupling between DMN connectivity and alpha power. (**A**) Mean changes (Post – Pre) in the dynamic coupling matrix for the Active (upper right) and Sham control (lower left) groups across subjects. (**B&C**) Boxplots illustrate increased coupling (from Pre to Post) between fluctuation of alpha power and PCC-rAG (**B**) and PCC-lAG (C) connectivity in the Active (vs. Sham) group. The red and blue shaded areas correspond to the mean ± 1.96 × SEM and the mean ± 1.96 × SD, respectively. For the double contrasts, * = *p* < 0.05 FDR corrected, ·= *p* < 0.1 FDR corrected. For the follow-up simple contrasts, · = *p* < 0.1, * = *p* < 0.05.

**Figure 3.**
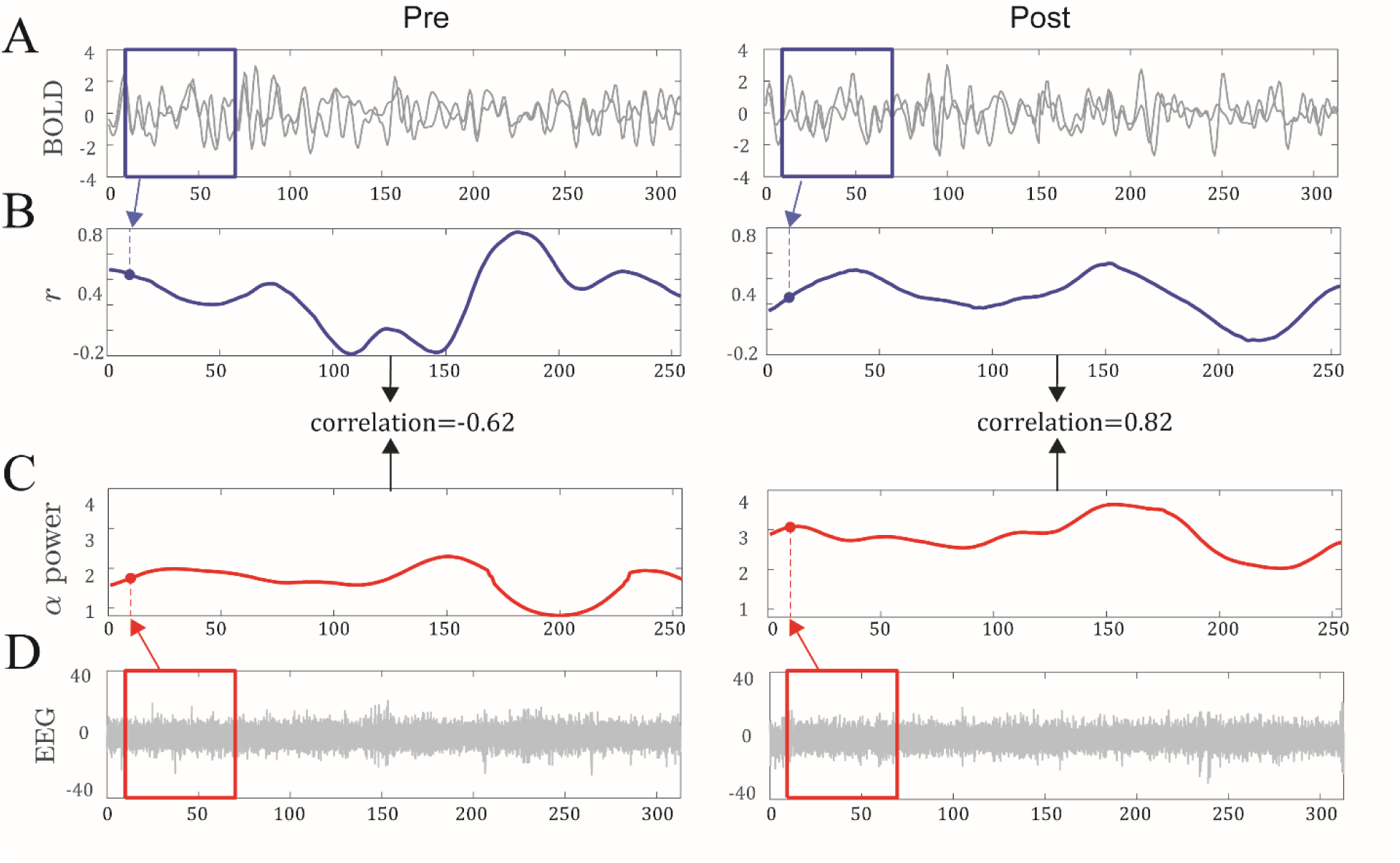
Data from a representative participant. (**A&B**) For each pair of ROIs (here, vPPC and right AG), average BOLD signals were extracted using a sliding Gaussian window of 60 TRs (∼2 minutes; **A**). The two BOLD timeseries for each sliding window were then correlated, resulting in a *r* coefficient (i.e., FC strength). Congregating the *r* values for all windows (sliding in increments of 1 TR) across the Pre (Left) and Post (Right) recordings, we obtained the respective FC timeseries (**B**). (**C&D**) Similarly, we obtained an alpha power timeseries for the Pre (Left) and Post (Right) sessions (**C**). Specifically, we extracted alpha power from each sliding window and congregated alpha power from all windows for Pre and Post sessions, repsectively (**D**). Finally, we correlated the timeseries of FC and alpha power for each session (**B & C**) and obtained an index of dynamic alpha-FC coupling. In this participant, the coupling increased from -0.62 Pre (Left) to 0.82 Post (Right).

### Effects were specific to the alpha frequency and DMN

To ascertain whether the coupling was specific to the alpha frequency, we performed similar tests on theta and beta frequencies. We observed no effect of α-tACS on these couplings (FDR *p*’s > 0.2; **Figure 4A**).

**Figure 4.**
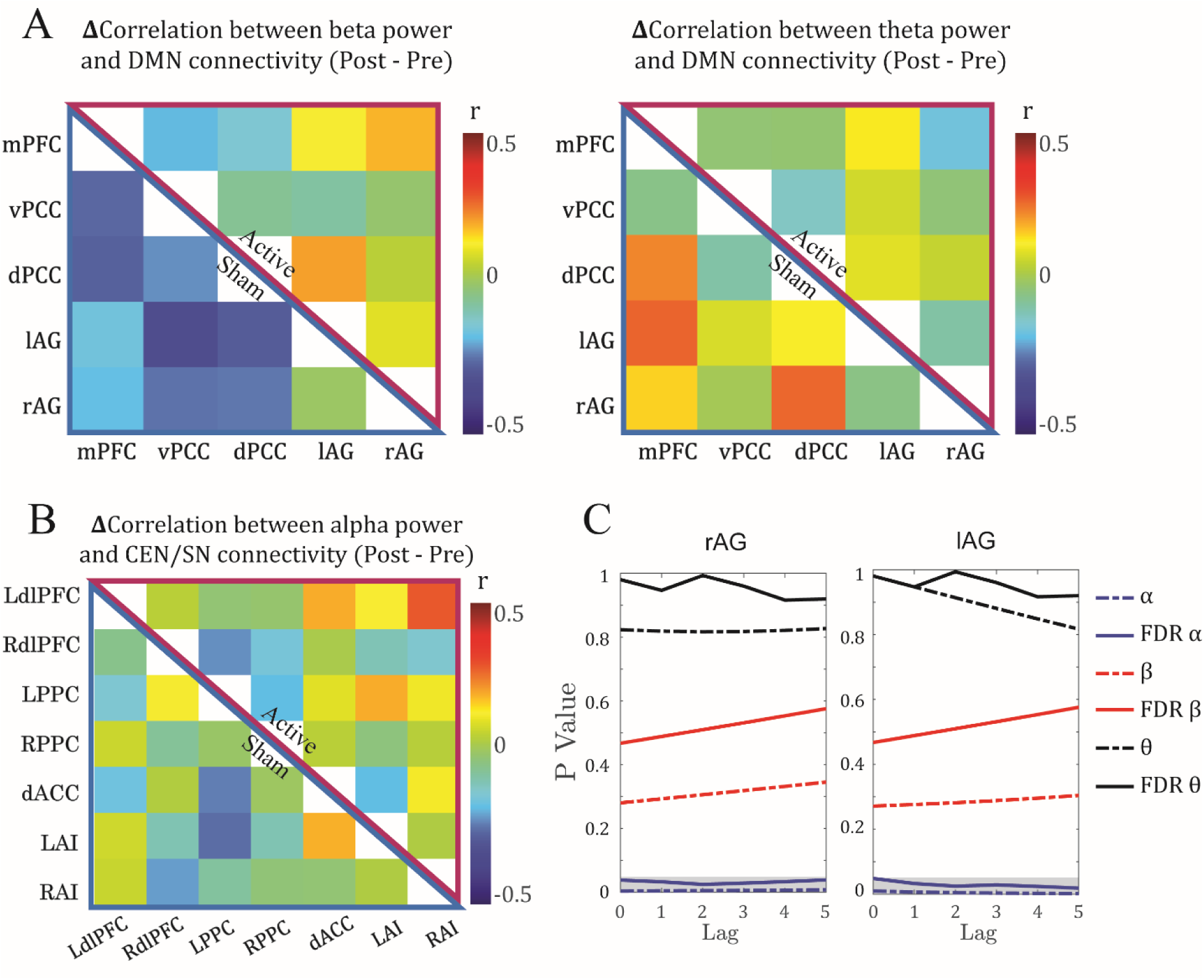
*α*-tACS modulation of dynamic coupling across brain networks and frequency bands. (**A**) No effect of α-tACS on the dynamic coupling between DMN connectivity and beta or theta power. Mean differential (Post – Pre) dynamic coupling matrices were comparable between Active (Upper right) and Sham (Lower left) groups for beta (**Left**) and theta (**Right**) band power. (**B**) No effect of *α*-tACS on the dynamic coupling between alpha power and CEN/SN connectivity. Mean differential (Post – Pre) dynamic coupling matrix of connectivity within and between CEN and SN was comparable between Active (Upper right) and Sham (Lower left) groups. No effects survived the significance threshold (*p* < 0.05 FDR corrected). (**C**) Effects of lag size. P- and corrected P-values for the effect of tACS on the dynamic coupling of alpha (blue lines) with vPCC-rAG (Left) and vPCC-lAG (Right) connectivity were consistently significant (grey shaded area)over lags of zero to five TRs. In contrast, beta (red lines) and theta (black lines) showed stable, non-significant patterns over the same lags.

To evaluate whether the coupling was specific to the DMN, we performed similar double contrasts on connectivity of the other major networks—the CEN and SN. As illustrated in **Figure 4B**, there were no effects of tACS on these couplings (FDR *p* ’s > 0.09).

### Effects were consistent across different lags

Finally, we examined the potential effect of lag size on the alpha-DMN dynamic coupling to consider the sluggishness of fMRI relative to EEG indices for neural activity [51, 52]. We varied the lag from 0 to 5 TRs (1.8 to 9 seconds), and as illustrated in **Figure 4C**, the results for vPCC-rAG and vPCC-lAG remained significant across the range (FDR *p* ’s < 0.05).

### Control Analysis

#### Controlling for alpha association with global connectivity fluctuations

To highlight the dynamic coupling of alpha with intra-DMN connectivity and control for global changes in DMN connectivity, we extracted the timeseries of extra-DMN connectivity between any of the DMN nodes with any of the CEN and SN nodes. We then subtracted the averaged correlation coefficients between alpha and extra-DMN connectivity timeseries from the corresponding correlation coefficients between alpha and intra-DMN connectivity and submitted the differential (i.e., adjusted) coefficients to the double contrasts for hypothesis testing.

We confirmed increase in dynamic coupling between alpha power and the adjusted vPCC-rAG FC in the Active (versus Sham) group (*t* = 3.8, FDR *p* = 0.006) after stimulation (**Figure 5**). Specifically, the Active group showed an increase (*t* = 3.35, *p* = 0.002, one-tailed), while the Sham group showed a decrease (*t* = -1.91, *p* = 0.038 one-tailed) in this coupling. We also found an increase in coupling between fluctuations of alpha power and the adjusted vPCC-lAG connectivity in the Active (versus Sham) group (*t* = 2.75, FDR *p* = 0.033) after stimulation.

**Figure 5.**
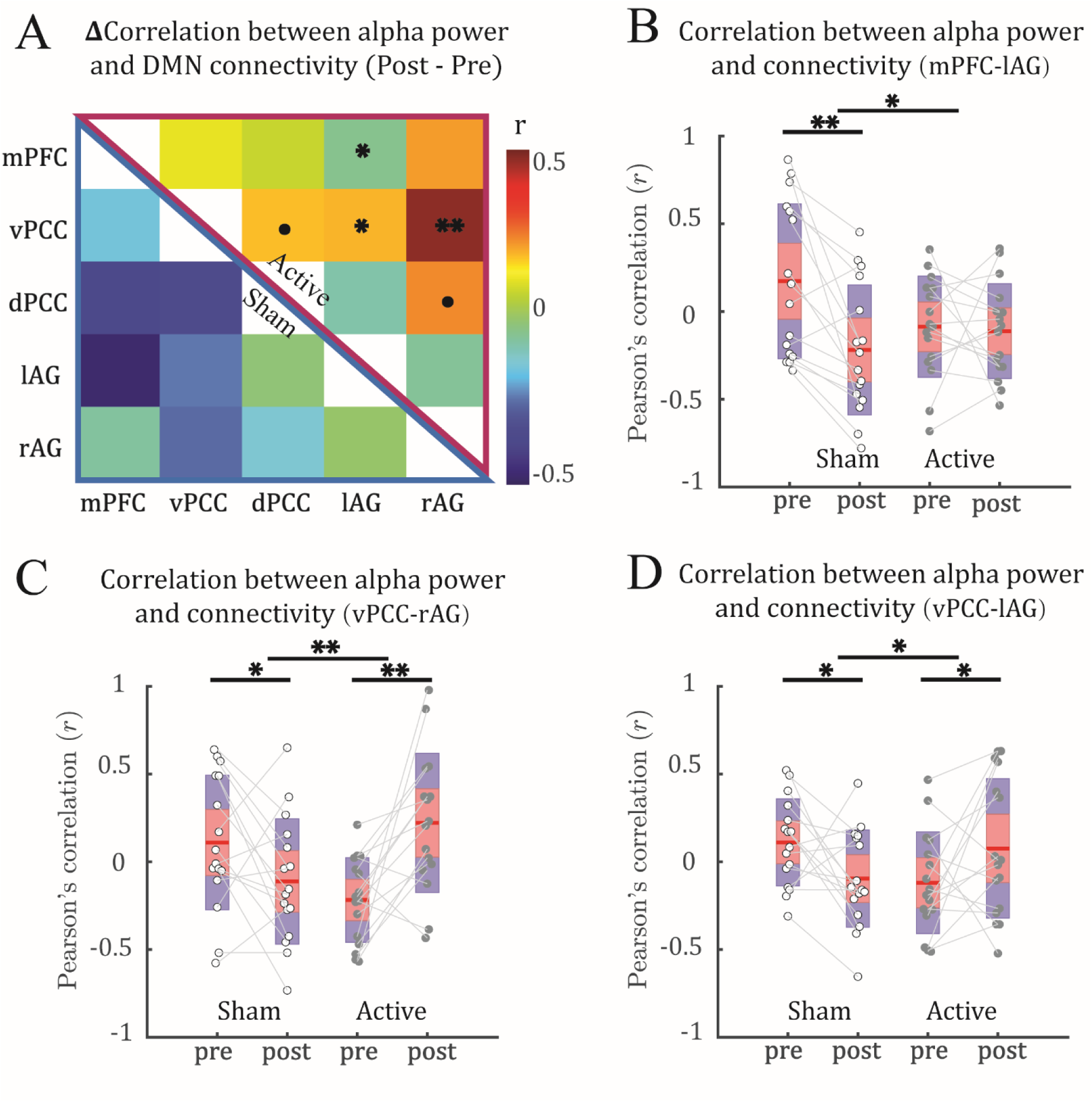
*α*-tACS strengthened dynamic coupling between adjusted DMN connectivity and alpha power. (**A**) Changes (Post – Pre) in the dynamic coupling matrix for the Active (upper right) and Sham control (lower left) groups. (**B-D**) Boxplots illustrate increased coupling (from Pre to Post) between fluctuation of alpha power with mPFC-lAG connectivity (**B**), PCC-rAG connectivity (**C**) and PCC-lAG connectivity (**D**) in the Active (vs. Sham) group. The red and blue shaded areas correspond to the mean ± 1.96×SEM and the mean ± 1.96×SD, respectively. For the double contrasts, · = *p* < 0.1 FDR corrected, * = *p* < 0.05 FDR corrected. ** = *p* < 0.01 FDR corrected. For the follow-up simple contrasts, · = *p* < 0.1 uncorrected; * = *p* < 0.05 uncorrected; ** = *p* < 0.01 uncorrected.

Specifically, the Active group showed an increase (*t* = 1.75, *p* = 0.049, one-tailed) while the Sham group showed a decrease (*t* = -2.19, *p* = 0.022, one-tailed) in this coupling. Moreover, we found an increase in coupling between fluctuations of alpha power and mPFC-lAG connectivity in the Active (versus Sham) group (*t* = 2.81, FDR *p* = 0.033) after stimulation. Specifically, the Active group showed no difference (*t* = -0.27, *p* = 0.39, one-tailed) while the Sham group showed a decrease (*t* = -3.96, *p* < 0.001, one-tailed) in this coupling.

#### Between-network connectivity effects

For completeness, we also examined the effects of tACS on EEG-fMRI coupling for between-network connectivity. tACS effects on the adjusted DMN connectivity were not observed for connectivity between DMN and CEN/SN (*p’s* > 0.05 FDR corrected). Notably, there was no effect of tACS for connectivity between the posterior DMN (PCC, r/l AG) and the posterior CEN (r/l PPC). A detailed picture illustration is available in the extended data (Extended Data Fig. 1-1).

#### Baseline alpha-DMN coupling

Pooling Pre-stimulation baseline data from the Active and Sham groups, we examined the baseline coupling between alpha and DMN connectivity timeseries. However, there was no reliable coupling between them (*p*’s > 0.1). A detailed picture illustration is available in the extended data (Extended Data Fig. 1-2).

## Discussion

Alpha oscillations and the DMN represent the dominant activities of resting-state EEG/MEG and fMRI, respectively. They are thought to play crucial roles in the spatiotemporal organization of the complex brain system, both figuring importantly in fundamental cognitive processes and implicated, often concurrently, in diverse neuropsychiatric disorders. Leveraging simultaneous EEG-fMRI data acquired before and after α-tACS, we demonstrated tightening of the coupling between temporal fluctuations in alpha power and DMN connectivity (particularly, in the posterior DMN between PCC and bilateral AG). Importantly, we systematically ruled out the effect of α-tACS on the coupling between fluctuations in beta and theta oscillations and in other major large-scale networks (the CEN and SN). In addition, findings were consistent across different time lags between fMRI and EEG timeseries, indicating the robustness of this alpha-DMN coupling. Therefore, the perturbation of alpha oscillations via α-tACS caused specific and robust impacts on the alpha-DMN coupling, highlighting an inherent association between these two major intrinsic neural activities. Moreover, the efficacy of α-tACS in enhancing this association provides support for tACS-based therapeutics in clinical interventions.

Functional connectivity and neural oscillations both fluctuate over time, and such dynamics bear significant behavioral and clinical relevance [23, 53, 54]. Accordingly, establishing a specific coupling between fluctuations in alpha and DMN connectivity would support an inherent association between alpha oscillations and the DMN. However, to date, only a handful of studies have leveraged simultaneous EEG-fMRI to perform direct investigation of such a coupling [17, 49]. Moreover, while this literature has demonstrated a negative coupling between fluctuations in resting-state alpha oscillations and between-network (e.g., DMN-CEN) connectivity, it has failed to uncover a reliable coupling between alpha power and DMN connectivity. We suspect that this coupling may be obscured by the various artifacts inherent in resting-state recordings, necessitating more powerful methods for its detection. α-tACS is an effective tool to modulate alpha oscillations [55, 56], experimentally perturbing its fluctuations and putatively, its coupling with the DMN. Indeed, by combining it with simultaneous EEG-fMRI, we garnered evidence for the alpha-DMN-connectivity coupling.

Interestingly, this coupling appears largely confined to the posterior DMN (between the PCC and the AG). Given that α-tACS was administered at the posterior scalp, targeting the occipitoparietal cortex, it is possible that this effect in the posterior DMN merely arose from a posterior alpha entrainment by α-tACS. However, α-tACS did not produce such an effect for connectivity between the posterior DMN nodes (PCC, left/right ANG) and the posterior node of the CEN (left/right PPC), thereby ruling out this explanation. While both are integral hubs of the DMN, the mPFC and PCC exhibit opposite connectivity patterns with other networks and show opposite responses during tasks, suggesting that they dissociable subdivisions of the DMN [4, 57]. Functionally, the PCC (and the posterior DMN) is believed to perform broad-based continuous sampling of external and internal environments [57]. Relatedly, alpha oscillations play a significant role in modulating sensory processing and maintaining vigilance [1, 2]. Therefore, there may be a special intimacy between alpha oscillations and the posterior DMN, which are inherently coupled to synergistically facilitate and regulate sensory sampling and vigilance. Sensory processing and vigilance were not examined in the current study to confirm this implication. However, previous research [31] has demonstrated the effect of α-tACS in modulating sensory processing and reducing anxious arousal, lending support to this notion.

Further in line with this idea, alpha oscillations and the DMN may share a fundamental anatomical substrate—the thalamocortical circuitry [58]. This circuitry has long been known to generate and regulate alpha oscillations [59, 60]. Recently, it has also been considered as a key controller of the DMN [4]. The thalamocortical circuitry, encompassing the thalamus, sensory cortex, and thalamic reticular nucleus, is a primary machinery for basic sensory processing. In addition, it is crucial for the control and regulation of arousal and vigilance [61]. Therefore, the coupling between alpha and DMN activity can arise from the thalamocortical circuitry to facilitate cooperation in sensory sampling and vigilance. Moreover, the effect of psilocybin, a serotonin agonist [62], can increase both alpha power and posterior DMN connectivity, implicating a neurochemical (serotoninergic) basis for the alpha-DMN coupling. Particularly, the serotonergic system plays a pivotal role in sensory processing and vigilance [63, 64], reinforcing the idea that alpha and DMN are inherently coupled to maintain vigilance in concert with environmental stimuli. Consequently, as the levels of sensory sampling and vigilance spontaneously wax and wane over time, alpha and DMN activity in resting-state recordings rises and falls in synchrony, manifesting as a tight temporal coupling.

The study has several limitations that need to be addressed in future research. First, as illustrated in **Figure 2**, the Active and Sham groups differed in baseline levels of alpha-DMN coupling. This could be a random sampling error as a result of our modest sample size (*N* = 32). That said, this difference may not present a major confound and obscure our hypothesis as the two groups showed opposite changes after stimulation, with the Sham group showing a decline in coupling overtime while the Active group exhibited an increase. Namely, tACS could overcome a general decline overtime and increase the alpha-DMN coupling. Second, potential confounding factors driving both DMN connectivity and alpha power include respiration and arousal, which naturally occur during rest and are often treated as noise in fMRI studies [65–68]. However, these fluctuations may have neural correlates, as suggested by significant correlations observed between alpha EEG power, respiration, and BOLD signals [69, 70]. Future research should incorporate specific measurements of such physiological variations to more specifically elucidate the relationship between alpha power and DMN connectivity.

In conclusion, by combining simultaneous EEG-fMRI timeseries and neural perturbation via α-tACS, we identified a specific temporal coupling between alpha oscillations and DMN connectivity. This finding lends further credence to an inherent, potentially mechanistic relationship between alpha oscillations and the DMN. Notably, the isolation of the posterior DMN in this coupling raises an intriguing possibility that this alpha-DMN coupling could bear particular relevance to sensory vigilance. Clinically, the efficacy of α-tACS in tightening alpha-DMN coupling presents new opportunities to noninvasively enhance neural functioning and treat mental disorders, especially those involving hypervigilance and sensory anomalies, such as posttraumatic stress disorder.

## Acknowledgments

This research was supported by the National Institutes of Health grants (R01MH132209 and R01NS129059 W.L.).

## Extended Data

**Figure 1-1.**
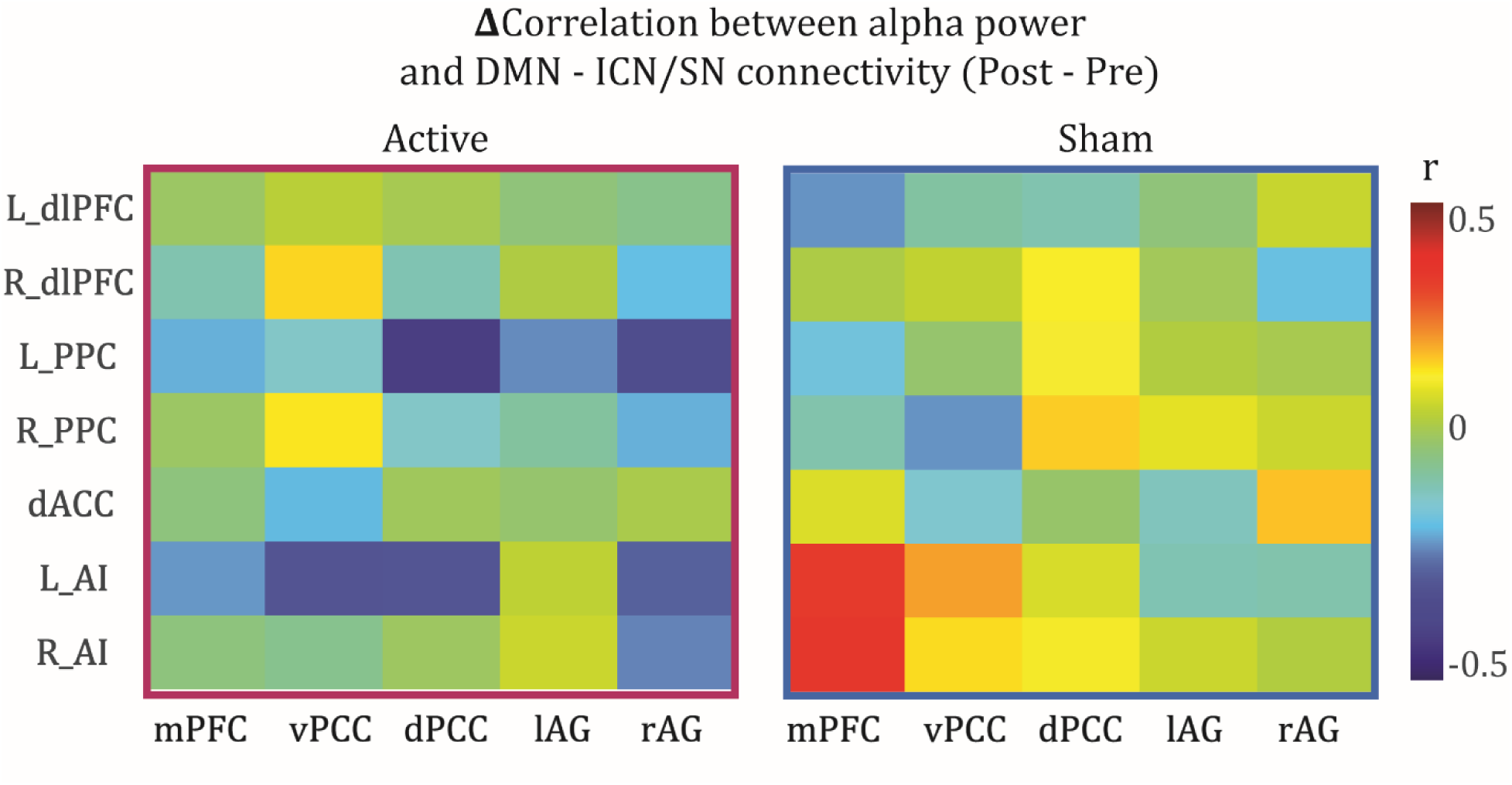
No effect of *α*-tACS on the dynamic coupling between alpha power and connectivity of DMN with the CEN and SN: Differential (Post – Pre) dynamic coupling matrices were comparable between Active and Sham groups. No significant effects of tACS were observed.

**Figure 1-2.**
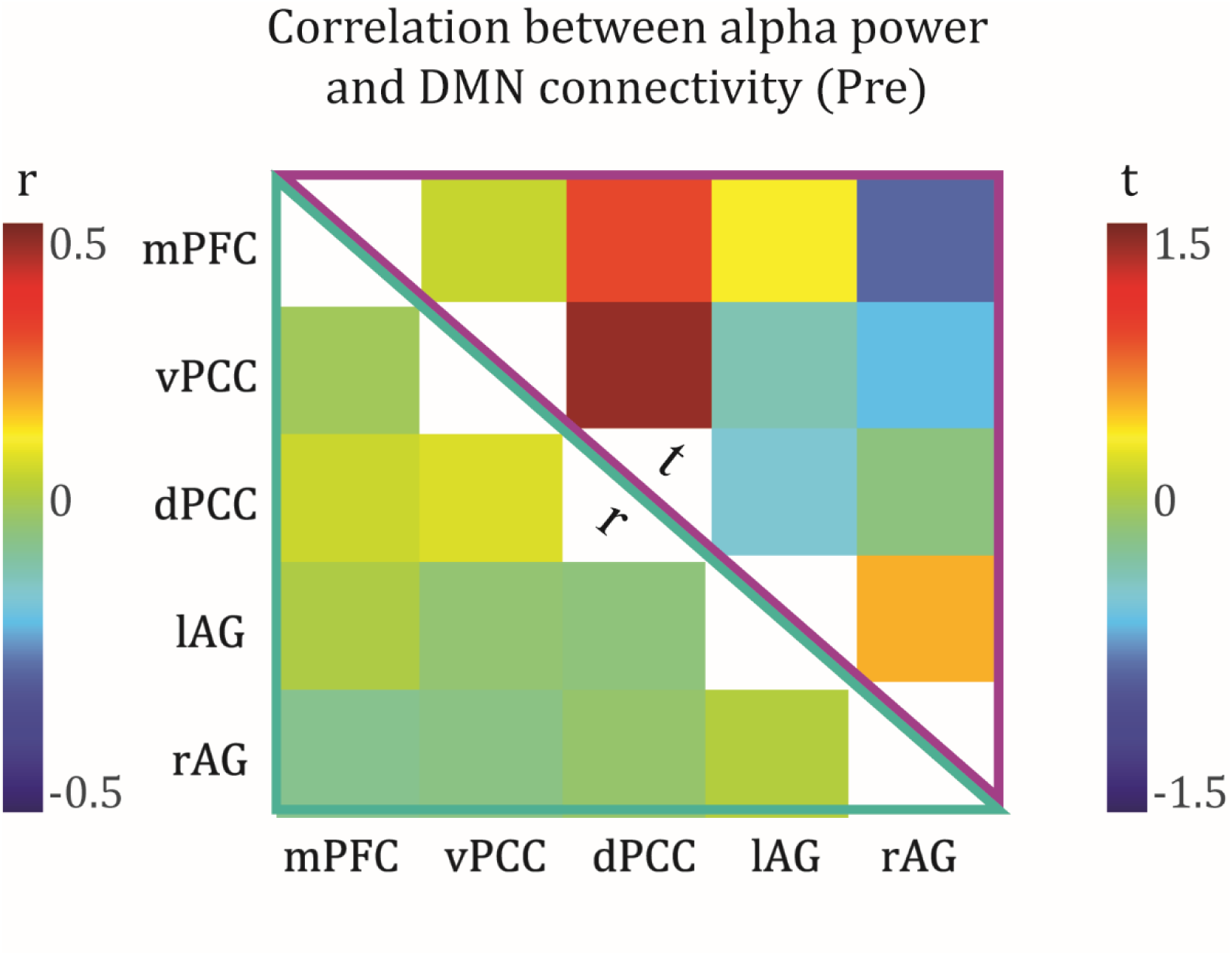
Baseline alpha-DMN Coupling: Baseline data pooled between the two groups showed no reliable coupling between alpha and DMN connectivity timeseries. The lower left and upper right portions of the matrix reflect the r and t values, respectively.

